# Sensory convergence in the world’s largest cavefish diversification: patterns of neuromast evolution, distribution and associated behaviour in *Sinocyclocheilus*

**DOI:** 10.1101/2021.12.10.472160

**Authors:** Bing Chen, Tingru Mao, Yewei Liu, Wenzhang Dai, Xianglin Li, Amrapali P. Rajput, Jian Yang, Joshua B. Gross, Madhava Meegaskumbura

## Abstract

The genus *Sinocyclocheilus* represents the largest freshwater cavefish genus in the world. This emerging model system is endemic to the southern Chinese karstic landscape, and demonstrates multiple adaptations for caves (troglomorphism), with eye-degeneration being the most pronounced. The less-apparent lateral line system, which is often expanded in cave-dwellers, has been studied in other cavefish systems, but never in the context of this diversification. Here we investigated the distribution and evolution of cephalic neuromasts in 26 *Sinocyclocheilus* species. We used live-staining and behavioural assays, and interpreted results in a phylogenetic context. We show that asymmetry in neuromast features and the rate of evolution is greater in cave-adapted species. Ancestral state reconstructions show that most *Sinocyclocheilus* are right-biased with some scatter, and show convergence of neuromast phenotypes. There is substantial variation in cephalic neuromast distribution patterns between and (to a lesser extent) within species. Behavioural assays show blind species have a distinctive wall-following behaviour. We explain these patterns in the context of the deep evolutionary history associated with this karstic region, other organismal traits, and habitat occupation of these remarkable diversifications of fishes. Interestingly, some of these neuromast patterns and behaviour show convergence with other phylogenetically distant cavefish systems.

## Background

Cave-dwelling (hypogean) fish provide a valuable system to study evolutionary novelty [1-4], owing to the extreme conditions associated with these habitats, such as limited food and perpetual darkness. Strong selective pressures arising from this extreme environment are associated with a suite of regressive morphological changes, such as loss of eyes and pigment [5, 6]. However, numerous constructive features also evolve in cave-dwellers. Among the most notable are expansions of the mechanosensory lateral line system [7]. Here, we examine the evolution of the lateral line system in the species-rich genus *Sinocyclocheilus*, an emerging model system and the largest diversification of freshwater cavefish in the world [8, 9].

Neuromast organs are composed of hair cells with cilia embedded in a gelatinous cupula, surrounded by a complex set of support cells [10, 11]. There are two general neuromast types: canal neuromasts (CN), which are larger, and embedded in a bony canal, and superficial neuromasts (SN), which project directly from the skin epithelium [12-14]. In the Mexican tetra (*Astyanax*), cave morphs harbor variation in the number and size of both types of neuromasts in the lateral line, including several-fold more SNs compared to surface fish. Moreover, *Astyanax* cave morphs have a highly asymmetric distribution of neuromasts across the left/right axis compared to surface-dwelling morphs [15]. At present, it is unclear the relevance of this asymmetry, however, some studies suggest it facilitates lateral swimming preference, rheotaxis (orientation towards flow), foraging, shoaling, predator avoidance, and mate-finding [7, 16-20].

*Sinocyclocheilus* are distributed across a vast 62,000 km^2^ karstic area in Yunnan and Guizhou Provinces and Guangxi Zhuang Autonomous Region [21, 22]. Based on their habitat preference, these fish can be divided into three principal groups: troglobitic (cave-restricted); troglophilic (cave-associated); and surface-dwelling [9]. However, one study characterized neuromast size differences between two *Sinocyclocheilus* species [20]. *Sinocyclocheilus* species span the macroevolutionary transition from surface to cave-restricted species and thus offer an exceptional opportunity to examine the evolution of traits associated with this transition.

Here we examined numerical variation, lateral distribution and behavioural differences associated with neuromasts in *Sinocyclocheilus*. Neuromasts may provide functional compensation for eye loss, therefore we hypothesized that more severe eye/vision loss may be associated with compensatory expansions of the lateral line neuromasts. Specifically, troglomorphy was predicted to be associated with more, and larger, neuromasts. Additionally, we anticipated a higher prevalence of distributional asymmetry of neuromasts in troglomorphic species compared to surface species, owing to relaxation of symmetry.

We found that neuromast distributions were asymmetric among all groups, however the degree of asymmetry was most apparent for Regress-Eyed (troglomorphic; collectively both Blind and Micro-Eyed) species. Additionally, in phylogenetic contexts the pace of neuromast evolution is faster among species with smaller eyes, compared to normal eyed species. Taken together, this work provides the first diversification-wide description of lateral line evolution, and clarifies the shared evolutionary pressure on constructive trait evolution among distantly-related species that colonize caves.

## 2. Methods

*Sinocyclocheilus* adult fishes used in this study were examined using live, biosafe staining techniques and behavioural assays. The project was approved by Guangxi Autonomous Region Government and Guangxi University Ethical Clearance Committee (# GXU-2021-125).

### (a) Fish maintenance in captivity

Adult fish used in this study were collected from the field from 2017-2020 across Yunnan and Guizhou Provinces and Guangxi Zhuang Autonomous Region of China (figure 1, see geographic coordinates in electronic supplementary material: table S1, figure S5). All fish were maintained in a centralized fish aquarium system, and maintained at (pH: 7.0 - 8.0, temperature 19°C ± 1°C, dissolved oxygen 8.5mg/L). Specimens larger than 5 cm were maintained in large glass aquaria (90 × 50 × 50 cm, 150 × 80 × 80 cm), with individual mechanical and bio-filters. For this study, 76 individuals from 26 *Sinocyclocheilus* species were used (table 1). *Carassius auratus* (n = 3) and *Cyprinus carpio* (n = 3), two species from a closely related clade to *Sinocyclocheilus* for comparison.

**Table 1.** Summary of 26 *Sinocyclocheilus* species used in the current analysis grouped according to eyes morphology and habitat.

|  | **Troglobitic (TB)** | **Troglophilic (TP)** | **Surface (S)** |
| --- | --- | --- | --- |
| **Blind (B)** | *S. furcodorsalis* |  |  |
|  | *S. tianlinensis* |  |  |
|  | *S. tianeensis* |  |  |
|  | *S. xunlensis* |  |  |
| **Micro-eyed (ME)** | *S. mashanensis* |  |  |
|  | *S. altishoulderus* |  |  |
|  | *S. microphthalmus* | *S. multipunctatus* |  |
|  | *S. bicornutus* |  |  |
|  | *S. rhinocerosus* |  |  |
|  | *S. cyphotergous* |  |  |
| **Normal-eyed (NE)** | *S. guilinensis* |  |  |
|  | *S. huanjiangensis* |  |  |
|  | *S. macrophthalmus* |  | *S. angustiporus* |
|  | *S. guanyangensis* |  | *S. oxycephalus* |
|  | *S. brevis* | *S. longibarbatus* | *S. malacopterus* |
|  | *S. lingyunensis* |  | *S. purpureus* |
|  | *S. zhenfengensis* |  | *S. maitianheensis* |
|  | *S. jii* |  |  |
|  | *S. yishanensis* |  |  |

**Table 2.** Model-averaged rate parameters for the measured traits of neuromasts in eyes-related morphs of *Sinocyclocheilus* species. **Bold font** = highest rate

| Trait | Average rate |  |  |
| --- | --- | --- | --- |
|  | Blind | Micro-Eyed | Normal-Eyed |
| Number | 155.1042 | **179.4815** | 150.6202 |
| Left Number | **0.6395** | 0.4106 | 0.4249 |
| Right Number | **0.7095** | 0.5056 | 0.5329 |
| Area | 102.8836 | **106.2819** | 101.9520 |
| Mean distance | **34.4129** | 11.7357 | 7.9001 |
| Density | **362.2959** | 64.3977 | 42.2944 |

**Figure 1.**
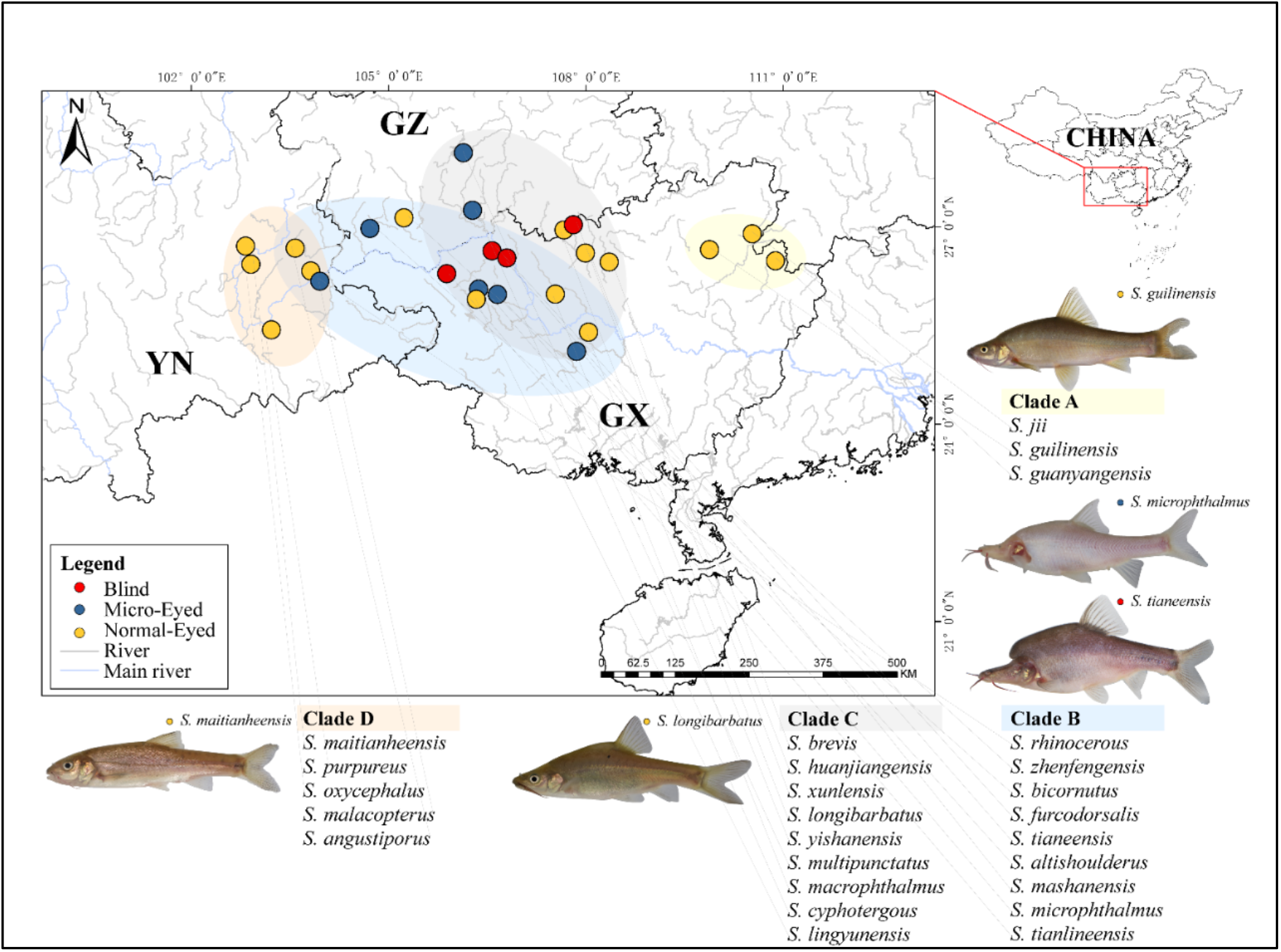
Map showing the distribution of sampling sites of the 26 species of *Sinocyclocheilus* (n = 76) used in this study. These fishes are mainly distributed across South China’s Karstic habitats on three provinces: Guangxi Autonomous region (GX), Guizhou (GZ) and Yunnan (YN). The four main clades (A – D, represented by a color code), the species used from each clade, and a representative photo of species from these clades are shown. Early emerging Clade A is mainly distributed in the eastern range of the diversification and Clade D to the hills to the West; Clade B and C contain the most troglomorhpic species and are distributed mainly across GX and GZ. The eye-condition (Blind, Micro-eyed and Normal-eyed) are depicted using a color code.

### (b) Vital staining of neuromasts

To visualize bones and neuromasts, live fish were immersed in 1mg/L Calcein-AM (C0875-5G; Sigma-Aldrich) and 20μg/ml DASPEI (D0815; Sigma-Aldrich) [15, 23, 24]. Fish were anesthetized using MS222 (E808894-5g; Macklin) 0.01-0.02g/L for 3-6 min followed by brief immersion in ice water for ∼20s, depending on size and age (e.g., smaller specimens were exposed to ice water for shorter periods). All individuals were continuously monitored to ensure the health and safety of experimental individuals.

### (c) Digital Analysis of Neuromast Position and between Distances

This study focused on the cephalic region anterior to the opercular and interopercular bones, near the arc on the lateral aspect [20, 22]. Images were collected using a Leica M165FC fluorescent stereomicroscope outfitted with Leica Application Imaging software (LAS X v3.4.2.18368). The dorsal, left and right aspects of each individual were imaged under identical conditions. High-definition “montage” image were consolidated following automated alignment and ‘flattening’ of 30 images collected in the z-plane (× 9/ 13/ 16 magnification) using LASX or Helicon Focus (Pro v7.6.1), to perform the two-dimensional shape images. The use of montage imaging minimized potentially confounding variables arising from the z-plane for each specimen.

The outline of fish and neuromast positions were obtained using the “Pencil” and “Point” tool in GIMP (v2.10.24). Neuromast numbers were quantified using an automated method in ImageJ (v1.8.0), following the method of Gross et al. (2016) [15]. For ambiguous images, we performed direct, manual counting (figure 3*a*). We calculated the size (2-D area) of individual neuromasts, lens, and eye orbit diameter by determining their size in pixels using the “measure” function in ImageJ and converting to mm^2^. We also used a vernier caliper to measure standard length (SL). To determine the density of neuromast within particular unit areas, we used “Delaunay Voronoi” triangulation in ImageJ to define a proxy for the mean distance between neuromasts [25-27] (figure 3*a*).

### (d) Quantifying symmetry

To examine positional symmetry of neuromasts across the left-right axis, we manually superimposed the fluorescent images of neuromasts (figure 3*b*). We used excitation filters for three colors (Blue 470/40 nm, Texas Red 560/40 nm and Green 546/10 nm). In GIMP, we then reflected the left images to align with the right images, creating a single image. We measured the scoring of neuromasts in the reverse sequence (right-side first) relative to the initial scoring of neuromasts (left-side first) to avoid bias in our calculations [28].

The “colocalization” [29, 30], and “JACop” [31, 32] plugins in FIJI were used to calculate an asymmetry coefficient [15, 33, 34]. We performed a Pearson’s correlation to compare the intensity distribution between channels [35]. Next, we calculated an Overlap Coefficient (OC), to identify positional overlap of signals from the left and right sides [36]. This enabled us to quantify the extent to which positions of neuromasts on the left and right sides of the cranium were symmetric (figure 3*a*).

We divided 26 species into Regressed-Eyed groups (containing Blind: n = 12 and Micro-Eyed: n = 21) and Normal-Eyed groups (n = 45), following the categorization and phylogeny of Mao et al. (2021). Importantly, a few *Sinocyclocheilus* species do not have uniform eye sizes [22]. For instance, *S. guanyangensis* have highly variable eye sizes, so for this study we selected individuals with the most common Normal-Eyed morphology. All parametric neuromast measures were subjected to one-way ANOVA with a post-hoc Tukey test analysis. Non-parametric distributions were subjected to Wilcoxon Signed Rank Test (2-tailed, Holm correction) or a Kruskal-Wallis test with a post-hoc analysis Dunnett test (2-tailed, Bonferroni correction). Statistical significance was set at p<0.05. We used the packages “wmc” and “FSA” in R (v3.6.3). Principal Component Analysis (PCA) was used to evaluate the following variables: number, area, asymmetry coefficients and mean distance. To analyse the relationships between Regressed-Eyed and Normal-Eyed groups, we performed Spearman’s rank correlation coefficients using SL, eye traits, area, number, mean distance and OC of neuromasts patterns.

### (e) Phylogenetic inference

To study the patterns of neuromast evolution in *Sinocyclocheilus*, we inferred a phylogenetic tree for the 26 species, using two available gene fragments from Genbank (NADH-ubiquinone oxidoreductase chain 4 - ND4 and cytochrome b gene – Cyt b), together with an outgroup species (*Cyprinus carpio*) (table S1). We used two different methods for phylogenetic inference [37] – Bayesian and Maximum Likelihood for tree construction (alignment, model selection and phylogenetic inference method details are available in the electronic supplementary material, Supplementary Methods).

### (f) Evolution rate analysis

To test the allometric evolution rate between neuromasts in different morphs [9], we analysed neuromast numbers, left/right-side numbers, mean distance coefficient, area and density (i.e. the proportion of number and area anterior to the gill). We used 100 potential trait histories from stochastic character mapping, and fit two alternative models (single or multiple rate model, calculated by AIC) of evolution for each studied trait. In the case of small samples, we assumed the AIC and AICc to assess, and then weighted from Bayesian analyses on the trees using the brownie.lite function. We used the package “rgl”, “ape” and “phytools” in R to calculate the model-averaged estimates [38].

### (g) Patterns of neuromasts evolution using ancestral state reconstructions

To understand the broad patterns of neuromast evolution in *Sinocyclocheilus*, we classified 26 species into the following three morphological groups: (1) Left-right axis asymmetry: according to different degree of overlapping coefficient, divided as Absolute-asymmetry (OC < 0.1) and Slight-asymmetry (OC ≥ 0.1) based on Gross et al. (2016) [15]. (2) Left/right-bias handedness: we used the normalized SN number by signed (directional) asymmetry rate: 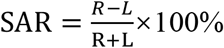 following Planidin et al. (2021) [39], which suggests the presence of two morphological categories: Right-biased (SAR > 0, neuromasts on right-side mainly) and Left-biased (SAR < 0, figure S1). (3) Distance expansion: depending on the results of neuromasts mean distance coefficient, we divided species into Scattered (DEL > 0.2) and Serried (DEL ≤ 0.2), following Gross et al. (2016) [15] (table S1).

To understand the evolution of neuromast patterns, we performed ancestral state reconstructions using these three morphological categories. One hundred stochastic reconstructions were simulated through the stochastic mapping approach that was conducted using function “make.simmap” (model = “ER”) in the R packages “ape”, “phytools” and “viridisLite”.

### (h) Cave-dwelling behaviour associated with neuromast variation

To understand how neuromast patterns may facilitate certain behaviour, we carried out a series of behavioural assays. All assays were performed using the following 14 species (n = 3 individuals for each): Blind - *S. furcodorsalis, S. tianlinensis, S. tianeensis*; Micro-Eyed - *S. mashanensis, S. microphthalmus, S. bicornutus, S. multipunctatus*; Normal-Eyed - *S. guilinensis, S. longibarbatus, S. macrophthalmus, S. oxycephalus, S. zhenfengensis, S. purpureus, S. maitianheensis*. Each experimental fish was acclimatized for 30 min in a rectangle assay arena (45 × 28 × 28 cm) in system water. An infrared camera (Cannon XF400/405) was used to capture video under infrared light in a quiet, dimly lit room (frame rate: 4Mbps (VBR, YCC 4:2:0, 25p, 1280 × 720).

Wall-following behaviour has been observed in *A. mexicanus* cavefishes; therefore, we examined a range of wall-following within the arena. This included fish swimming a minimum distance of its SL along the wall, or a distance of ≤ 0.5 SLs away from the wall (figure 5*a-b*) [40, 41]. We recorded normal tracking without stimulation for 10-min to determine if wall-following behaviour was present using EthoVision XT (v15.0, Noldus) alongside direct visual monitoring. (2) We next used an aeration pump (45∼50 Hz vibration) for 5-min to record the approaching of a novel object in modified vibration attraction behaviour test [42-44]. We selected the area centered on the stimulation model based on tank shape (10 × 16 cm). We monitored the frequency of time spent in the stimulation range from left/right-side. Data analysis were performed in R’s basic functions as mentioned.

## Results and Discussion

Neuromast distribution is asymmetric in all cave fish studied to date [15, 45-48]. Accordingly, we found the two outgroup species of Cyprinidae showed several neuromasts on observation region, which did not show a comparable pattern of neuromast asymmetry as was observed in *Sinocyclocheilus*. Fish living in perpetual darkness frequently lose vision while enhancing non-visual sensation, such as the lateral line [42, 49, 50]. Here, for the first time, we show that *Sinocyclocheilus* similarly conform to this pattern (figure 2). Additionally, widespread neuromast distributional asymmetry in *Sinocyclocheilus* is convergent with other distantly related cavefish species inhabiting similar cave microenvironments. Interestingly, our results showed that this asymmetric neuromast pattern is variable across the 26 *Sinocyclocheilus* species tested, with most species demonstrating a right-sided bias (figure 4). Evidence for neuromast asymmetry patterns in other lineages comes from two lineages from Central and North America: *Astyanax mexicanus*, with a left-side bias and the *Gasterosteus aculeatus* (stickleback fish), with a right-side bias [39, 51]. These convergent results reveal variation in “sidedness” across taxa, despite convergence in the asymmetry of neuromasts traits.

**Figure 2.**
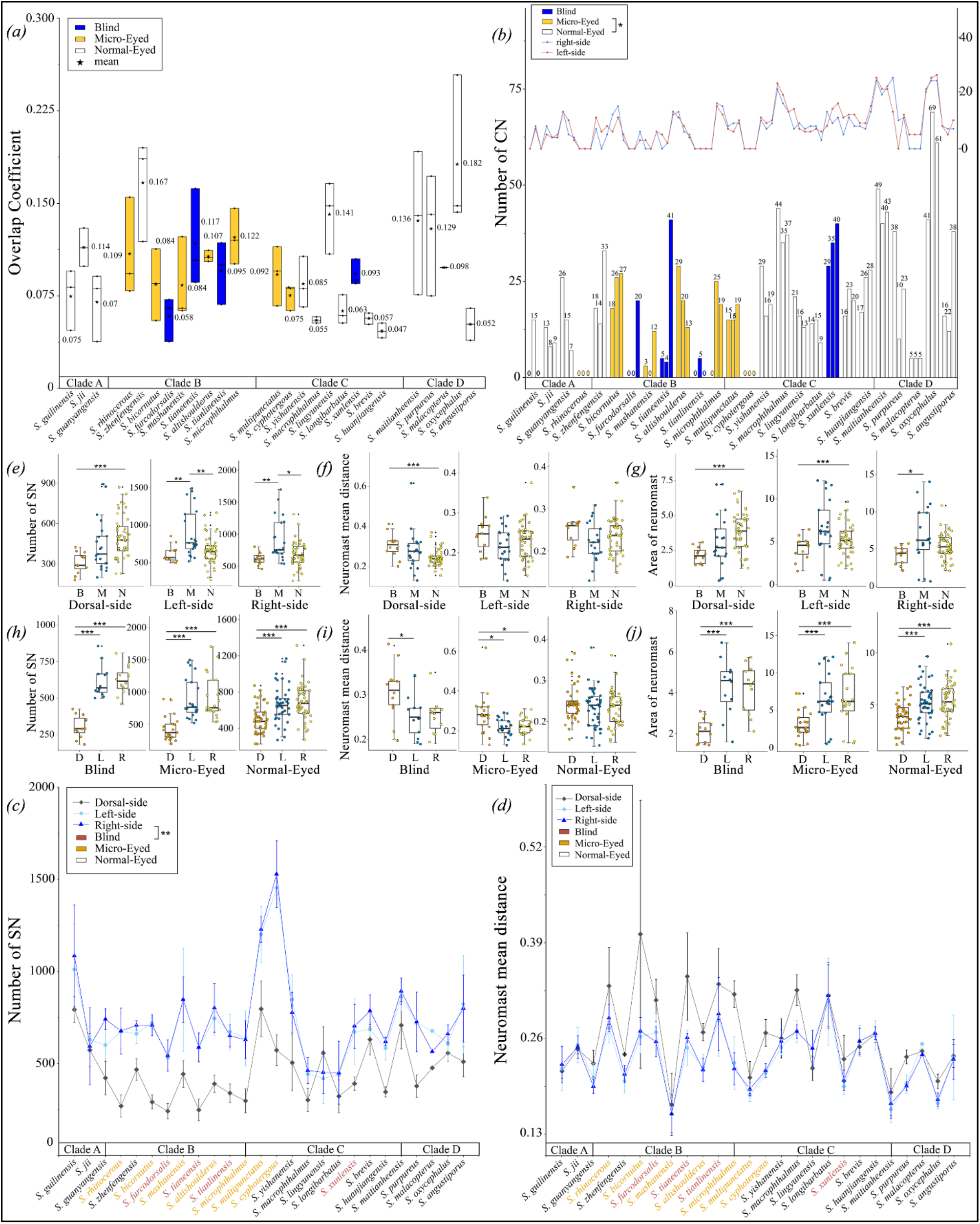
Comparison of neuromast-related measurements for 26 species of *Sinocyclocheilus* (n = 76). (a) The average scores of Overlapping Coefficient (OC) measured as the pattern of neuromasts on left/right-side of the fish. The standard of asymmetry results is defined as: asymmetry (OC < 0.1), symmetry (OC ≥ 0.1). Clade A-D represent the clade level relationships of these 26 species in the phylogenetic tree. All statistical results are available in table S1, S2. (b) The bars represent the individual’s canal neuromasts (CN) total counts and the lines represent the left and right sides of cephalic CN in different species. The CN of Micro-Eyed species was significantly less than Normal-Eyed species (H = -2.64, p = 0.025). (c) The mean number of superficial neuromasts (SN) on the cephalic area (dorsal, left and right sides) in different *Sinocyclocheilus* species. (d) The mean surface area covered by SN on the dorsal/left/right-side of different *Sinocyclocheilus* species. The surface species covered by the dorsal SN are less than that of the surface are covered by the right and left sides of fish. (e-g). The comparisons about: (e) the number of SN; (f) neuromasts mean distance coefficient; (g) area of SN on Dorsal/Left/Right-side on cephalic area. Group divided by the Blind (B in orange), Micro-Eyed (M in blue) and Normal-Eyed (N in yellow). The Wilcoxon signed rank (2-tailed, Holm correction) test suggested that the surface covered by dorsal neuromasts are significantly less than that covered by the right and left sides. ^*^: P < 0.05, ^**^: P < 0.01, and ^***^: P < 0.001. All statistical results are available in table S2, S5. (h-j). The comparisons of the non-parametric statistical tests performed for (h), the mean number of SN; (i) neuromasts mean distance coefficient; (j) mean area of SN on the cephalic area of the Blind, Micro-Eyed and Normal-Eyed morphs. The comparisons were sub-divided into dorsal-side (D in orange), left-side (L in blue) and right-side (R in yellow) for visualization purposes.

### (a) Patterns of neuromast symmetry and asymmetry

The neuromasts distribution pattern showed asymmetry across the left-right axis, with variation both within and across species (figure 2*a*, table S2). Interestingly, mean overlap coefficients indicated that all species showed a degree of neuromast asymmetry. Normal-Eyed species showed the least asymmetry, while Blind species showed the highest and Micro-eyed species had intermediate values (mean ± s.d.: Normal-Eyed = 0.098 ± 0.05; Blind = 0.091 ± 0.031; Micro-eyed = 0.096 ± 0.027; H_2_ = 0.64, p > 0.05). This finding is consistent with prior work suggesting that asymmetry may facilitate navigation in darkness [52, 53], foraging [54] and the ability to maximize sensory information using fewer receptors [39, 51]. In comparing asymmetry measures across clades, Clade D (all Normal-Eyed, surface-dwelling species, showed the least asymmetry (mean ± s.d. = 0.119 ± 0.058), while Clade C (mean ± s.d. = 0.079 ± 0.031) showed the highest asymmetry; with Clade A and B demonstrating intermediate values (Clade A – 0.086 ± 0.029 and Clade B – 0.104 ± 0.038).

### (b) Patterns of neuromast distribution

Generally, we found more SNs on the right compared to the left-side, however we observed substantial variation between and across species. In the dorsal aspect, Blind fish have the fewest, Normal-Eyed fish have the most, and the Micro-Eyed group was intermediate (figure 2*e*). Unexpectedly, there were significantly fewer SNs in Blind species (mean ± s.d.: Blind = 1543 ± 248; Micro-Eyed = 2252 ± 802; Normal-Eyed = 1859 ± 504; H_2_ = 10, p = 0.0065, table S5). In terms of lateral distribution, Blind and Normal eyed fish had higher numbers on the right-side, while Micro-Eyed species had more neuromasts on the left-side (median: left = 767, right = 758; W = 227, p = 0.890; figure 2*h*; table S5).

We found that dorsal neuromast numbers are significantly lower than the lateral sides. An interesting exception was *S. lingyunensis* (N-TB, Clade C), dorsally distributed neuromasts outnumbered laterally-situated neuromasts (mean dorsal = 557, left = 419, right = 453; figure 2*c*, table S5). In the dorsal aspect, Normal-Eyed species had the most neuromasts, while the Blind group had the fewest. One potential explanation for this difference could be that obligate subterranean fishes experience relaxed selective pressure (e.g. aerial predation from birds) [21, 55], but perhaps dorsal neuromasts are necessary for navigation within caves [56].

With respect to neuromast expansion (scatter), we found that the most scattered SNs are dorsal, especially in Blind species (mean ± s.d.: Blind = 0.31 ± 0.06; Micro-Eyed = 0.28 ± 0.10; Normal-Eyed = 0.24 ± 0.04; H_2_ = 12, p = 0.002; figure 2*f*, table S5). Additionally, neuromasts tend to be more scattered on right-compared to the left-side across all three groups (median right/left Blind = 0.26, 0.25; Micro-Eyed = 0.22, 0.21; Normal-Eyed = 0.24, 0.23; figure 2*d*). Interestingly, the Blind cavefish group demonstrated the most scatter, while the Micro-Eyed group had the least scatter (figure 2*i*, table S5). The Mexican cavefish (*Astyanax mexicanus*) demonstrates more scatter in the surface forms compared to *Sinocyclocheilus* [15, 57], however this is likely due to the fact that *Astyanax* Blind forms have more neuromasts, and hence less scatter.

In the comparison of surface area covered by neuromasts, we found a significantly lower surface area populated by neuromast on the cephalic region dorsally, with the Blind group having the smallest area (mean ± s.d. Blind = 2.14 ± 0.59; Micro-Eyed = 3.13 ± 1.94; Normal-Eyed = 3.88 ± 1.33; H_2_ = 15.9, p = 0.0003; figure 2*j*). When comparing lateral sides, we found the total distributional area of neuromasts to be highly comparable within groups (median right/left-side Blind = 4.44, 4.6; Micro-Eyed = 6.19, 6.21; Normal-Eyed = 5.31, 5.14; table S5). Normal-Eyed species have the largest area of neuromast distribution, Micro-Eyed fish have an intermediate area, and the Blind group had the smallest area covered by (mean ± s.d. = 10.56 ± 2.95; Micro-Eyed = 16.44 ± 8.80; Normal-Eyed = 14.49 ± 4.59; F = 3.83, p=0.026; figure 2*g*). Although it has been shown that the neuromast complexity of the Blind species is greater than Normal-Eyed species [20], this however, has not been established at a diversification-wide scale.

Blind *Sinocyclocheilus* have fewer CNs compared to Normal-Eyed *Sinocyclocheilus* (median Blind = 5, Micro-Eyed = 13; Normal-Eyed = 17; table S5), reflecting the same pattern as SNs. For *Astyanax*, surface fish are invariant in terms of numbers, but cavefish are highly variable. Surprisingly, two species, *S. rhinocerous* and *S. cyphotergou* appeared to possess no CNs (both M-TB, figure 2*b*), suggesting the canal lateral line system may have regressed in this species. Interestingly, a similar phenomenon has been observed in Amblyopsid cavefish (Teleostei: Percopsiformes) [47] and in Threespine Sticklebacks (*Gasterosteus aculeatus*) [58].

In sum, neuromasts are generally reduced in number and area in Blind species, but not in distribution (expansion). Somewhat surprisingly, Micro-Eyed species had the most neuromasts, greater area, and least dense distribution of neuromasts. We propose that Blind species may have optimized adaptation to the subterranean biome by using fewer, but more complex, neuromasts. Blind *S. tianlinensis*, for instance, have SNs with greater diameters and more hair cells compared to Normal-eyed *S. macrophthalmus* [20].

### (c) Correlations between asymmetry and eye condition

We performed a Spearman Rank’s correlation of the highly loading variables for the Regressed-Eyed and Normal-Eyed. In the Regressed-Eyed group, eye measures demonstrated a significantly positive correlation with neuromasts number (ρ= 0.7, p < 0.001) and area (ρ= 0.7, p < 0.001), but a slightly negative correlation with mean distance between neuromasts (ρ= -0.3, p < 0.05). In the Normal-Eyed group, eye measures had a significantly positive correlation with neuromasts area (ρ= 0.4, p < 0.005) and mean distance (ρ= 0.4, p < 0.05), but significantly negative correlation with the results of asymmetry (ρ= -0.4, p < 0.001; figure 3*d,e*, table S4). Overall, the Regressed-Eyed group is more asymmetric compared to Normal-Eyed group.

**Figure 3.**
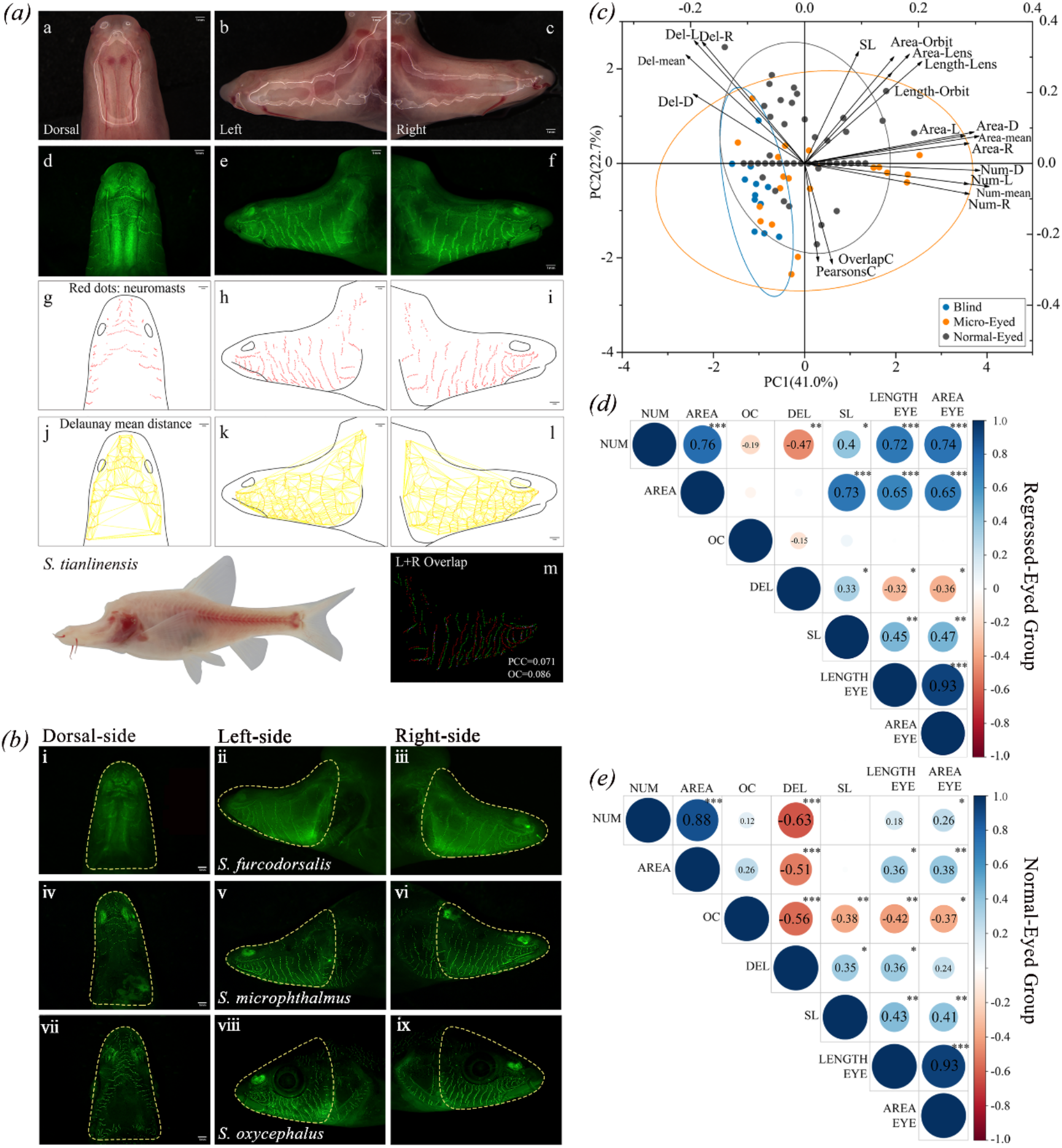
Summary of fluorescent staining results for different cavefish morphs. (a) the fluorescent staining results of *S. tianlinensis* (Blind species-Troglobitic, B-TB). (table 1 & table S1). (a-c) photos taken under normal lights. (d-f) dorsal, left and right sides of the neuromasts under fluorescent light. (small green dots are SN, bigger green dots are CN). (g-i) The diagrammatic representations of the dorsal, right and left side neuromasts. SNs denoted by red dots and the area outlined in black represent cephalic region and the olfactory area. (j-l) The neuromasts mean distance measures as the “Delaunay Mean Distance” are indicated by yellow lines. (m) The asymmetry values calculated by overlap coefficient. Red and green dots represent the left and right sides of the fish. Note that the scale of the images is the same at the length of 1.0 mm. (b) The results of the fluorescent staining of three cavefishes, (i-iii) *S. furcodorsalis*, (iv-vi) *S. microphthalmus* and (vii-ix) *S. oxycephalus*, used as representatives of each eye-related morphs. These three species are classified as (B-TB), Absolute-asymmetry/Scattered/Left-bias; (M-TB), Slight-asymmetry/ Scattered/Right-bias and (N-S), Slight-asymmetry/Serried/Right-bias respectively. Scales of the images are the same at the length of 1.0 mm. (c) Principal Component Analysis (PCA) bi-plot for the distribution traits of neuromasts in *Sinocyclocheilus*. The dorsal/left/right-side and mean counts number (Num-D, Num-L, Num-R, NUM); dorsal/left/right-side and mean distance coefficient (Del-D, Del-L, Del-R, DEL); asymmetry coefficients of neuromasts (PearsonsC, OverlapC); eyes traits (area of lens/orbit (Area-Lens, Area-Orbit) and length of lens/orbit (Length-Lens, Length-Orbit)) with standard length [5] were included as variables in the PCA. All results are available in Supplemental table S2, S3. (d) The Correlations between SN number (NUM), area (AREA), mean distances coefficient (DEL) and asymmetry coefficient (OC) of neuromasts; standard length [5] and the length of lens (LENGTHEYE) and area of lens (AREAEYE). The scores indicate the results of the spearman’s rank correlation coefficient |ρ|; < 0.3 = no correlation, 0.3-0.8 = low correlation, > 0.8 = high correlation. Positive correlations (ρ > 0) and negative correlations (ρ ≤ 0) are shown in blue and red colors respectively. ^*^: P < 0.05, ^**^: P < 0.01, and ^***^: P < 0.001 indicate the level of statistical significance. All results are available in table S4.

### (d) Dimensionality of the neuromast variables

We performed a principal component analysis (PCA) based on 26 species of *Sinocyclocheilus* and their neuromast-related variables, to identify key features underlying neuromast variation. Variation on PC1 was explained mostly by the number of neuromasts (mean number: NUM, counts on dorsal side: Num-D) and the total neuromast distributional area (AREA). PC2 was explained mostly by the distances between neuromasts (mean distance coefficient on left side: Del-L, on right side: Del-R; table S3). The 95% confidence intervals for each group shows that intraspecific variation in the Blind group was the narrowest, while the Micro-Eyed group showed the highest variation (figure 3*c*). This may be explained by the fact that Micro-Eyed forms are subjected to selective pressures of both surface habitats, subterranean habitats and the transitional habitats between these [22, 59, 60]; i.e. habitat heterogeneity.

### (e) Patterns of neuromasts evolution

We next performed an ancestral reconstruction for three character states: asymmetry, handedness and neuromasts expansion. We found the deeper nodes in the phylogeny showed ambiguity for asymmetry and handedness (see maximum credibility tree; figure 4). However, the deeper nodes for neuromast distribution suggested a scattered distribution of neuromasts is the ancestral condition. In *Sinocyclocheilus*, ∼ 70% of the species examined demonstrate right-handedness. Our ancestral state reconstruction shows an entire clade of 7 species is right-handed. However, both left- and right-handed mixed clades and sister taxa are present within the diversification, suggesting this is a variable trait. In contrast, ancestral state reconstructions show nearly 80% of the *Sinocyclocheilus* have scattered neuromast distributions. Additionally, all Blind species showed a scattered neuromast pattern of distribution. (figure 4*c*; black arrow).

**Figure 4.**
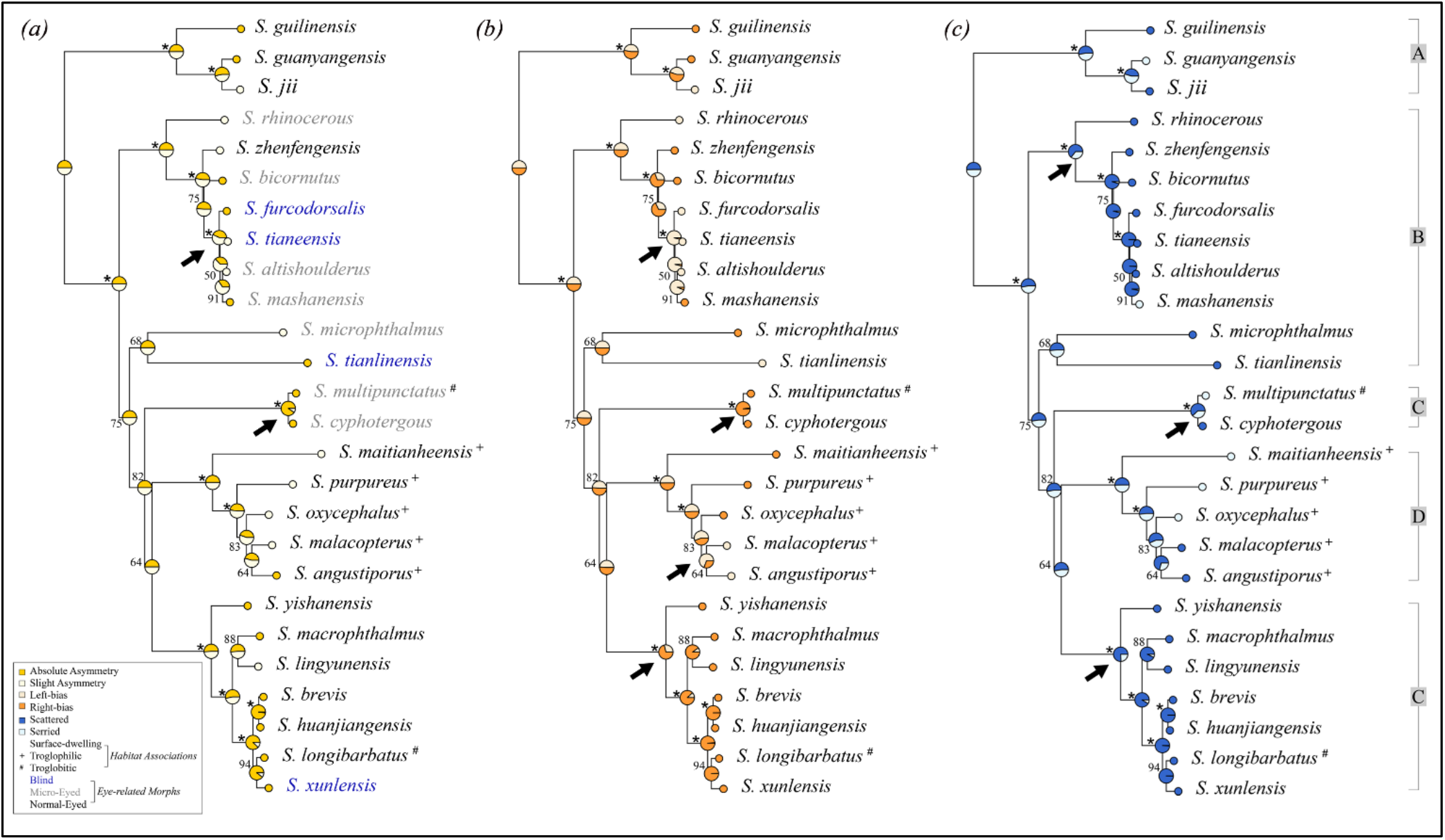
Patterns of cephalic neuromast trait evolution of *Sinocyclocheilus* based on ancestral state reconstruction on the maximum-likelihood tree. Ancestral state reconstructions based on the Bayesian inference method is shown in figure S2-S4. (a) Ancestral state reconstruction using stochastic character mapping for symmetry in neuromasts pattern (Absolute/Slight-asymmetry). The tip colors represent state of the extant species and each node indicates the ancestral state as a proportion of the tip state. A, B, C and D showed on the extreme right of the figure indicate four major clades. The bootstrap values > 95% are indicated as ^*^ on nodes. (b) Ancestral state reconstruction using stochastic character mapping for handedness bias (Left/Right-bias) on the phylogeny. (c) Ancestral state reconstruction using stochastic character mapping for neuromast expansion (Scattered/Serried) on the phylogeny.

### (f) Evolution rates analysis

Among cavefish, the rate of neuromast evolution has only been determined for *Sinocyclocheilus*. The rate of scatter for Blind species is higher compared to others (mean distance and density - 3.5 and 6.8 times increased than Normal-Eyed species), and the rate of numerical expansion of neuromasts is 1.2 times greater in Micro-Eyed species (table 3). Normal-Eyed *Sinocyclocheilus* species demonstrated lower evolutionary rates for every character state. A multiple-rate model of evolution provided the best fit for mean distance between neuromasts, relative neuromast density and neuromast distribution area. However, a single-rate model of evolution provided the best fit for traits associated with neuromast numbers (table S6). Thus, the evolutionary rate of cave-related neuromast patterns were highest in cave adapted forms. Further, rates of evolution for surface area covered by SN neuromasts, and right-sided neuromast numbers, are moderately increased in Blind and Micro-Eyed species.

### (g) Behavioural correlates of neuromasts

We performed a series of behavioural assays, and found that Blind species navigate markedly differently from eyed-species. Blind species have a well-established wall-following behaviour, while sighted species utilize an entire arena space (figure 5). We found that Blind species prefer to use the side with fewer neuromasts for wall-following, but the side with more neuromasts for exploring (i.e., stimulation during VAB test). Micro-Eyed species having a left-bias in neuromast distribution, and they preferred approaching the stimulation from the left-side (median = 4) more frequently than right-side (median = 3; W = 612, p>0.05). These species, however, followed the wall on their right-side (median = 154) for a significantly longer time compared to the left-side (median = 142; W = 967, p < 0.001; figure 5*c,g*; table S5). Behavioural heatmaps depicting wall-following behaviour showed that Blind species prefer swimming in a fixed route, but eyed-species swam in irregular patterns (figure 5 *d-f’*). Wall following behaviour has also been characterized in *Astyanax mexicanus*. In this species, blind morphs similarly use the side with fewer neuromasts towards the wall, perhaps to use the contralateral side for detecting stimuli important for feeding, communication and spatial learning [17, 42].

**Figure 5.**
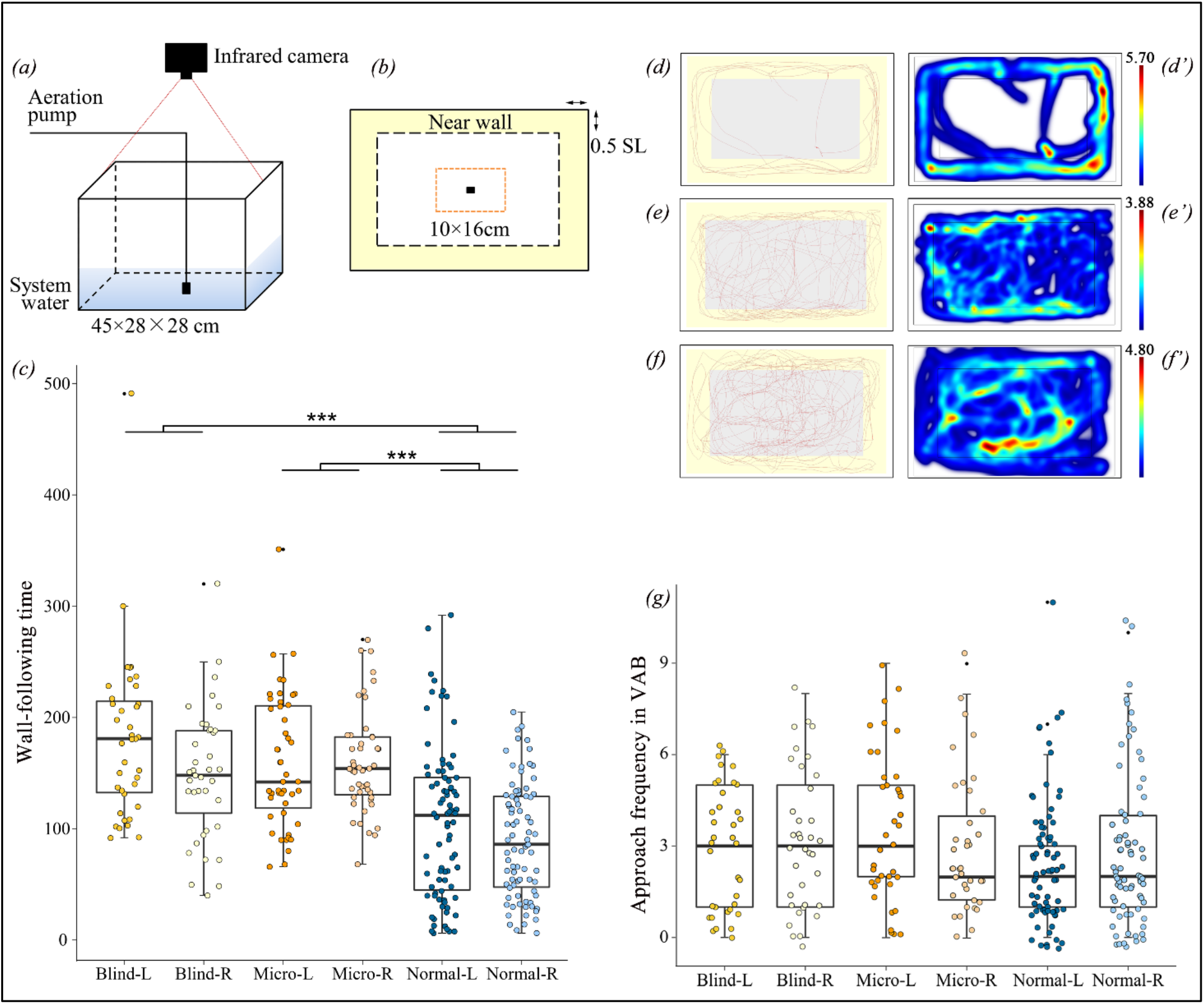
The experimental setup and results of the behavioural assays. (a) Schematic diagram showing the assay system used to for the wall-following and vibration attraction behaviour (VAB). (b) The vertical view of the arena’s schematic diagram. The pale yellow area within the dotted line depict the near-wall belt (“Near wall”). The orange dotted line showed the stimulating area. The black rectangle in center shows the Aeration pump. (c) The boxplot showing a comparison of the wall following time of Blind (3 species; n = 9), Micro-Eyed (4 species; n = 12) and Normal-Eyed (7 species; n = 21) morphs. Their handedness was also considered and the Kruskal-wallis Rank Sum Tests showed that the groups spent significantly different (U = 79.49, p < 0.001) times following the wall. Levels of significance indicated by ^***^. (d-f’). Representative result of a 10-min wall-following behaviour assay for three species. Vertical view of the swimming path of Blind fish (d,d’; *S. tianeensis*, B-TB), Micro-Eyed fish (e,e’; *S. microphthalmus*, M-TB) and Normal-Eyed fish (f,f’; *S. macrophthalmus*, N-TB). (d, e, f) Visualization of the traces of the fish swimming paths as depicted by EthoVision. (d’, e’, f’) Heatmaps generated from the trial results of the wall-following behaviour. The color bar represents the total time (min) the fish stayed in one place. Warmer colors denote areas with a longer time spent, whereas cooler colors denote areas of shorter time spent. (g) The results of the frequency of approach in stimulation area of VAB test.

*Sinocyclocheilus* is an ancient species complex, with an estimated age for the clade of nearly 10-11 Mya [9, 62].They are distributed across an enormous karstic area spanning three provinces in China [21]. One of the main forms of speciation seems to be isolation of populations over long periods, and therefore geographic speciation appears to have dominated this diversification [61]. This divergence occurred during the geological uplift of the Yunnan-Kweichow Plateau together with the aridification of China, which occurred during the Pliocene and the Pleistocene [61]. The neuromast variability seen in these fishes is most likely attributable to the collective influences of both selection- and drift-related evolutionary mechanisms that have played on these fishes over long periods. However, the exact dynamics of the evolution of neuromasts is still an intriguing question to be explored.

## Conclusion

In *Sinocyclocheilus*, alongside some basic patterns, there is widespread variation in cephalic neuromast patterns between species and to some extent within species. We showed neuromast asymmetry with right-side enhanced for most species. For almost all species, the dorsal neuromast numbers were lesser than either of the sides.

Furthermore, Regressed-Eyed (Blind and Micro-Eyed) species are more asymmetric than the Normal-Eyed forms. Interestingly, we found the greatest degree of neuromast variation and rate of evolution in Micro-Eyed species (living outside and inside caves - troglophilic), this is possibly an adaptation for life in two markedly different habitats types. Assays of swimming behaviour suggest a functional role of neuromasts in habitat exploration. These patterns of neuromast distribution and swimming behaviours are convergent with other cavefish lineages. The diversity of patterns and variation can be explained by the deep evolutionary history associated with the karstic region and the associated traits of this remarkable diversifications of fishes.

## Abbreviations Section

TB: Troglobitic
TP: Troglophilic
S: Surface
B: Blind
M: Micro-Eyed
N: Normal-Eyed
CN: Canal neuromast
SN: Superficial neuromast
GXU: Guangxi University
SL: Standard length
DEL: Delaunay Mean Distance (neuromast mean distance coefficient)
L: Left-Side
R: Right-Side
D: Dorsal-Side
PCC: asymmetry coefficient 1 = Pearson’s Correlation Coefficient
OC: asymmetry coefficient 2 = Overlap-Coefficient
PCA: Principal component analysis
AIC: Akaike information criterion
mtDNA: Mitochondrial DNA
ND4: NADH-ubiquinone oxidoreductase chain 4
Cyt b: cytochrome b gene.

## Declarations

## Acknowledgements

We thank the following individuals: members of EDD lab for their cooperation and support; Chenghai Fu for assistance in the field; Gajaba Ellepola for suggestions with the data analyses.

## Ethics approval

The project was approved by Guangxi Autonomous Region Government and Guangxi University Ethical Clearance Committee (protocol number: GXU-2021-125).

## Consent to participate

Not applicable.

## Availability of data and material

All the data used in the study are provided in the electronic supplementary material.

## Authors’ contributions

Conceptualization – MM, BC, JBG; Fieldwork – YL, BC, TM, MM, JY; Experimentation and Lab Work – BC, XL, APR; Data Analyses – BC, TM, WD, XL; Interpretation – All; Figures – BC, WD, TM; Writing original draft – BC, MM, JBG, APR; Subsequent Drafts – All; Funding acquisition – MM, JY, BC, TM, YL; Supervision – MM, JBG, JY.

